# Key residue Unlocks Dual PIP_2_-Dependent and Independent Gating in G Protein-Gated Inwardly Rectifying Potassium Channels

**DOI:** 10.1101/2025.02.14.637896

**Authors:** Ha Nguyen, Jonathan Mount, Keino Hutchinson, Yihan Zhao, Yulin Zhao, Ian W Glaaser, Peng Yuan, Avner Schlessinger, Paul A Slesinger

## Abstract

G protein-gated inwardly rectifying potassium (GIRK) channels mediate membrane hyperpolarization in response to G protein-coupled receptor activation and are critical for regulating neuronal excitability. The membrane phospholipid phosphatidylinositol 4,5-bisphosphate (PIP_2_) is essential for regulating a large family of ion channels, and disruptions in PIP_2_ interactions contribute to some neurological diseases. Structural analyses have identified key residues in PIP_2_-mediated gating of the GIRK2 channel. Notably, Arginine-92 (R92), a highly conserved basic residue at the membrane interface in GIRK2, interacts with PIP_2_ as well as cholesteryl hemisuccinate (CHS), a potentiator of GIRK2. Here, we used a combination of electrophysiological assays, fluorescent K^+^ flux measurements, cryo-electron microscopy, and molecular dynamics simulations, and find that mutations at R92 (Y, F, and Q) not only alter PIP_2_ sensitivity but can also reveal a novel gating mechanism that is independent of PIP_2_. These findings indicate that R92 plays a crucial role in modulating GIRK2 channel gating, offering new insights into developing potential therapeutic targets for treating neurological disorders linked to GIRK channel dysfunction.

## Introduction

G protein-gated inwardly rectifying potassium (GIRK) channels play a critical role in regulating the resting membrane potential and excitability of neurons^1,2^. When neurotransmitters bind to their cognate G protein-coupled receptors that link to pertussis toxin-sensitive G_i/o_ proteins, Gβγ subunits are liberated which directly bind to and activate GIRK channels, leading to membrane hyperpolarization and a decrease of neuronal excitability^3,4,5,6^. Of the four subtypes of GIRK channels, GIRK1, GIRK2, and GIRK3 isoforms are expressed predominantly in the brain^6,7,8^. GIRK1/2 heterotetramers form the most common GIRK channel in the brain, followed by GIRK2 homotetramers^1,8,9,10,11^. Research on mice and humans has implicated GIRK channels in several human diseases, including Down syndrome^12,13,14^, Parkinson’s disease^8,15,16^, Keppen-Lubinsky syndrome^17,18^, depression, and addiction^19^. A deficiency in GIRK channel activity is associated with an increased risk of seizures, alcoholism, and heightened sensitivity to psychostimulants^20,21,22,23,24^. Given their pivotal role in these neuropathologies, GIRK channels are a potential therapeutic target for treating neurological diseases^25^.

The interaction with the membrane lipid phosphatidylinositol 4,5-bisphosphate (PIP_2_) is crucial for the activation of all inwardly rectifying potassium (Kir) channels, including GIRK channels (also known as Kir3). The first evidence of direct PIP_2_ activation came from studies using excised membrane patches from *Xenopus* oocytes, where 50 μM of PIP_2_ was sufficient to activate GIRK1/4 and GIRK2 channels in the absence of exogenous Gβγ^26^. Conversely, the depletion of PIP_2_ through phospholipase C (PLC) activation decreases GIRK currents in patch-clamp electrophysiology experiments^26^. More recent findings using purified GIRK proteins reconstituted in liposomes have confirmed these results, demonstrating increased flux (decrease in fluorescence) from channel activation upon PIP_2_ application^27^. Additionally, several studies have shown that activators such as Gβγ G protein, cholesterol, and alcohol all enhance GIRK channel activity by increasing PIP_2_ relative affinity^27,28^. Overall, these studies have consistently established that PIP_2_ alone is sufficient to activate these channels.

X-ray and cryo-EM studies have shed light on how PIP_2_ interacts with GIRK channels at the structural level. These studies revealed a network of basic amino acids that stabilize PIP_2_ within its binding pocket on GIRK2. Specifically, the 1’-phosphate (PO_4_) group of PIP_2_ is coordinated by R92 in the first transmembrane helix (TM1) of GIRK2, while the 5’-phosphate (PO_4_) group interacts with positively charged residues K194, K199, and K200 in the second transmembrane helix (TM2)^29^. Additionally, K64 is positioned close to the anionic headgroup of PIP_2_^29,30,31^. PIP_2_ binding shifts the channel’s equilibrium between two conformations: an extended conformation, where the cytoplasmic domain (CTD) is disengaged from the transmembrane domain (TMD), and a docked conformation, where the CTD is engaged with the TMD^29,31^. PIP_2_ favors the docked conformation. Furthermore, structures of GIRK2 with both PIP_2_ and the cholesterol analog cholesteryl hemisuccinate (CHS) suggest that the docked state is further stabilized by these modulators^29^. Notably, in all resolved structures of GIRK2 with PIP_2_ and additional modulators such as CHS and Gβγ, the PIP_2_-binding site appears identical, offering limited insight into the mechanism of channel opening^29,30,31,32^.

Much of our understanding of PIP_2_ gating comes from mutagenesis studies, which reveal how changes in basic amino acids within the PIP_2_ binding site influence channel gating properties. For example, substituting the conserved positively charged K200 with tyrosine (K200Y) enhances PIP_2_ binding affinity^33^. Another study demonstrated that the K64Q mutation altered selectivity of GIRK2 for different phosphoinositides, favoring PI(4,5)P_2_ with specific acyl chains, while the K194A mutation increased preference for PI(3,4,5)P_3_ over other phosphoinositides^34^. These basic residues are critical not only for GIRK channel gating but also for other Kir channels. In Kir2.1 channels, replacing K182 (the equivalent of K194 in GIRK2) with glutamine weakened PIP_2_ interactions^35^. Additionally, mutating R82, K187, and K188 in Kir2.1 (corresponding to R92, K199, and K200 in GIRK2) significantly reduced whole-cell currents, underscoring the importance of these residues in maintaining channel functionality^35^.

Previous research has highlighted the importance of basic amino acids, such as lysine (K) and arginine (R), in coordinating the 4’ PO_4_ and 5’ PO_4_ of PIP_2_. However, less is known about the function of residues that interact with the 1’ PO_4_ of PIP2. In the cryo-EM structure of GIRK2 bound to PIP_2_ and cholesteryl hemisuccinate (CHS), R92 coordinates the 1’-PO_4_ of PIP_2_ while also forming a salt bridge with the CHS head group^29^. Despite this, the role of R92 in GIRK2 gating remains unclear. In this study, we investigate the role of the R92 residue in the PIP_2_-mediated gating of GIRK2 channels. We find that substitutions at position R92 result in GIRK2 channels that gate independently of PIP_2_. By combining electrophysiological data, fluorescent K^+^ flux data of purified channels reconstituted in liposomes, structural data obtained through cryo-electron microscopy (cryo-EM) single particle analysis, and MD simulations, we demonstrate that R92Y, R92F, and R92Q result in altered PIP_2_ sensitivity and evidence of K^+^ conduction in the absence of PIP_2_. Our findings offer structural explanations for the abnormal gating properties observed in R92 substituted channels and suggest an alternative gating mechanism associated with changes in this residue.

## Results

### R92 substitutions alter the alcohol and G-protein sensitivity of GIRK channels

To elucidate the role of R92 in gating of GIRK2 channels (**Figure 1A**, panel i), we substituted R92 with either alanine (A), tryptophan (W), phenylalanine (F), tyrosine (Y), lysine (K), or glutamine (Q) and assessed channel function in transiently transfected HEK293T cells co-expressing GABA_B1b_ and GABA_B2_ receptors (**Figure 1A**, panel ii). We measured the amplitude of the whole-cell inwardly-rectifying K^+^ current using the GIRK channel inhibitor BaCl_2_ (**Figure 1B** panel i-iv). Substituting R92 with non-polar residues (A, W, and F) led to a 50-60% reduction in basal GIRK2 currents, while mutation to polar or uncharged amino acid (Y and Q) resulted in substantial increases in Ba^2+^-sensitive basal currents. The conservative substitution of R92 with lysine (K), another positively charged amino acid, led to small basal currents (**Figure 1C**, panel i). Taken together, these results demonstrate that the positive charge at R92 is not essential for channel activity, and furthermore, substituting the charge with polar groups may enhance channel function.

**Figure 1.**
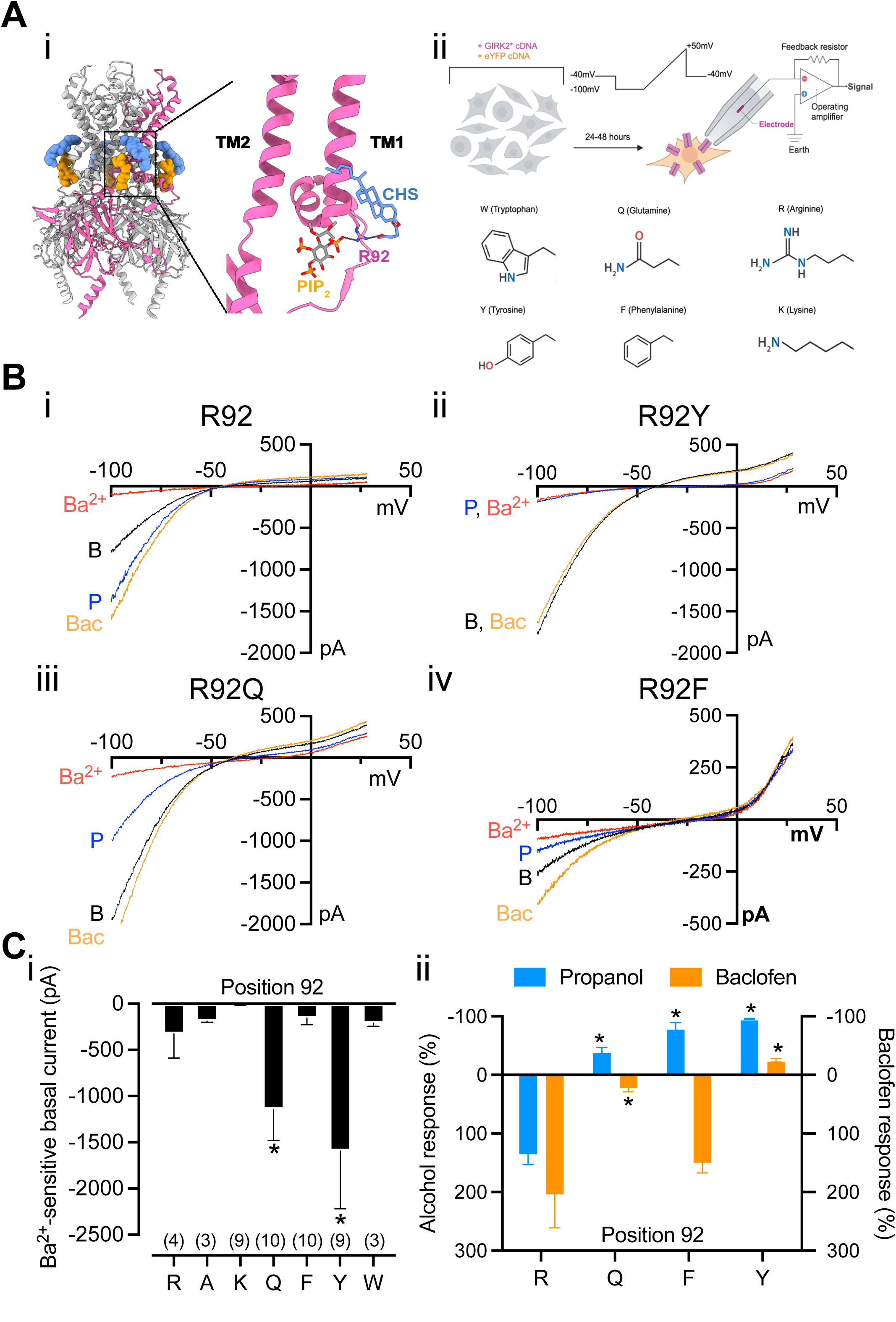
Structural and functional analysis of amino acid substitutions at R92 in GIRK2 channels. (**A**) A structural and experimental overview. (i) The GIRK2 channel structure (PDB:6XEV) highlights the position of R92 (magenta) near PIP_2_ (orange) and cholesteryl hemisuccinate (CHS, blue)^29^. The boxed region is magnified to show interactions at R92. (ii) Schematic of the electrophysiology measurements for R92 substitutions in GIRK2 channels expressed in HEK293 cells. R92 is substituted with W (tryptophan), Q (glutamine), Y (tyrosine), F (phenylalanine), or K (lysine), shown as corresponding side-chains. (**B**) Representative current-voltage (I-V) relationships are shown for R92 (i), R92Y (ii), R92Q (iii), and R92F (iv) channels in the presence of Ba²⁺ (red, 1 mM), Propanol (P, blue, 100 mM), Baclofen (Bac, orange, 100µM), or with 20K solution (B, black). (**C**) Quantitative analysis of substitutions at R92. (i) Bar graph shows amplitude of Ba²⁺-sensitive basal currents for the indicated R92 substitutions (*p < 0.05; ns, not significant). (ii) Percentage change from baseline with alcohol (Propanol, blue) or Baclofen (orange) for R92 and indicated substitutions (*p < 0.05).

To assess how mutations at R92 affect regulation of GIRK2 by alcohol and G proteins, we quantified the effects of 100 mM propanol and 100 µM baclofen, a GABA_B_ receptor agonist (**Figure 1B**, panel i-iv). For R92, propanol nearly doubled the basal current, consistent with literature reports of alcohol-induced GIRK activation^36^ (**Figure 1C**, panel ii). Baclofen also increased GIRK currents more than two-fold, indicating G protein-mediated activation (**Figure 1C**, panel ii). In R92Y, however, propanol inhibited the current to levels comparable to Ba^²⁺^ blockade. Baclofen also slightly inhibited the current with R92Y, in contrast to its activation of R92 channels. Similar inhibitory effects of alcohol were observed in R92F and R92Q, while baclofen induced slight activation in these channels, though to a lesser extent than in R92 channels (**Figure 1C**, panel ii). These results suggest that R92 substitutions retain basic channel functionality of conducting inwardly-rectifying K^+^ current, but the regulatory effects of alcohol and G proteins are significantly altered, underscoring the importance of R92 in GIRK channel activity. In addition, the significantly larger basal currents for R92Y and R92Q channels suggest some type of potentiation of channel function with these substitutions.

### R92 substitutions demonstrate PIP_2_-independent characteristics in K^+^ fluorescent flux assay

We hypothesized that the large basal currents observed in R92Y and R92Q channels could be attributed to enhanced PIP_2_ affinity. To test this, we co-expressed GIRK2 channels with Dr-Vsp, a voltage-dependent phosphatase that depletes membrane PIP_2_ by removing its 5′-phosphate, resulting in the inhibition of GIRK2 channels^37,38^ (**Figure 2A**). As expected, R92 channels were completely inhibited after 50-100 ms of Dr-Vsp activation (**Figure 2B,C**, panel i). On the other hand, R92Y, R92F, and R92Q channels exhibited a significant component of basal current that was resistant to PIP_2_ depletion, as indicated by the substantial K⁺ conductance retained even following prolonged Dr-Vsp activation up to 500 ms per pulse (**Figure 2B,C**, panel ii-iv). These results suggest that R92 mutations may either (1) confer high PIP_2_ affinity, preventing PIP_2_ from being removed by Dr-Vsp, (2) alter the phosphoinositide selectivity of the channel, allowing perhaps PI(4)P (a product of PIP_2_ dephosphorylation) to bind and activate the channels, or (3) enable a novel activation pathway that is independent of PIP_2_. Given that Kir2.1 channels, which have higher PIP_2_ affinity than GIRK2 channels, are fully inhibited by Dr-Vsp activation^33^, the first hypothesis seems unlikely.

**Figure 2.**
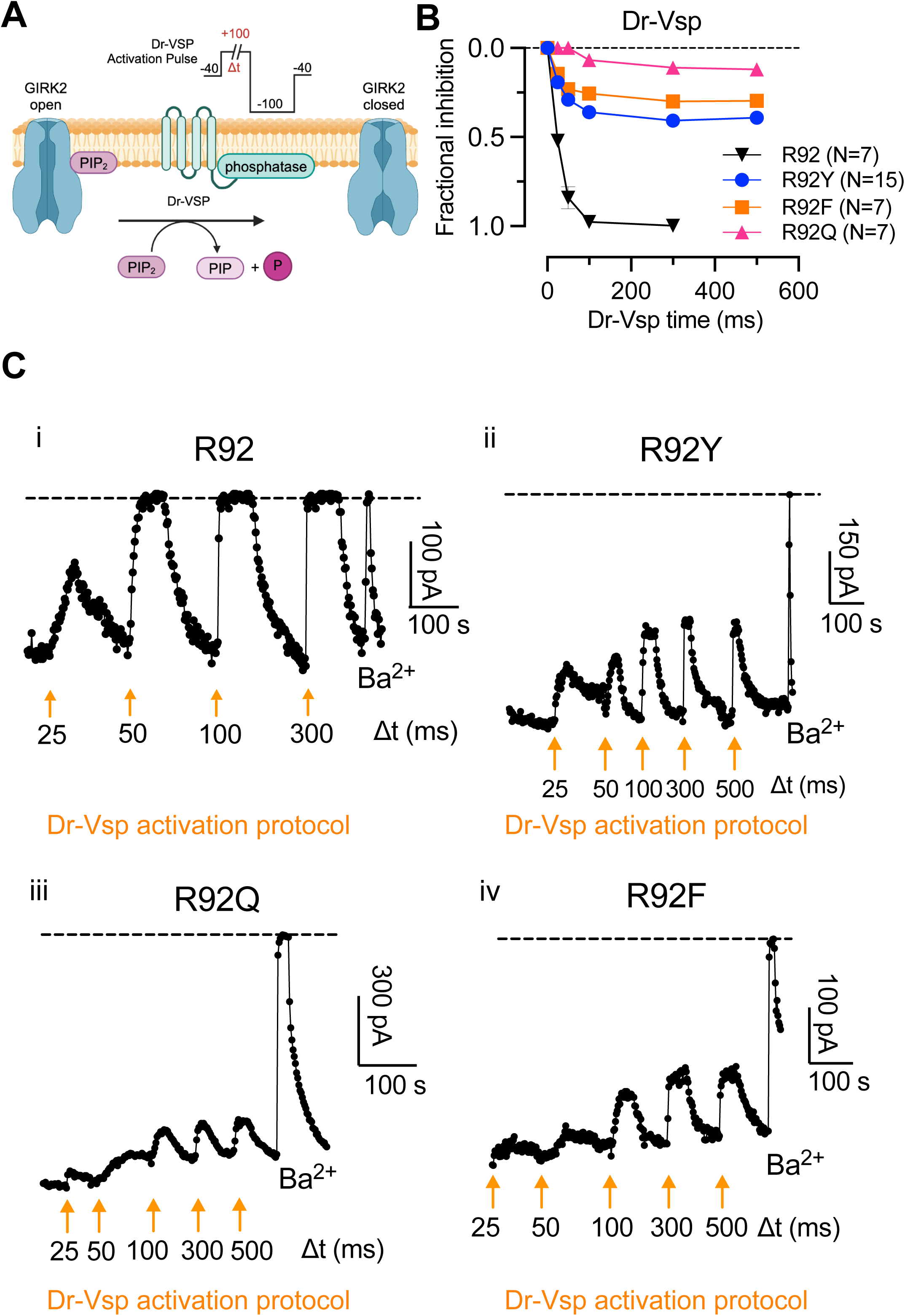
Effects of R92 substitutions on GIRK2 channel regulation by Dr-VSP-mediated PIP_2_ depletion. (**A**) Schematic of Dr-VSP-mediated PIP_2_ depletion. The protocol includes varying lengths (Δt) of depolarizing pulses to +100 mV. (**B**) Graph shows time-dependent inhibition of inward current by Dr-VSP activation for R92 and substitutions. Fractional inhibition of GIRK2 currents is plotted against time of Dr-VSP activation for R92 (black), R92Y (blue), R92F (red), and R92Q (pink) channels. Data are mean ± SEM (N indicates the number of cells). (**C**) Representative traces of GIRK2 currents during Dr-VSP activation. Time courses of current reduction are shown for (i) R92, (ii) R92Y, (iii) R92Q, and (iv) R92F in response to increasing lengths of Dr-VSP activation at +100 mV (orange arrows).

To further investigate whether altered phosphoinositide selectivity or a novel activation pathway explains the persistent activity in R92 substitutions following PIP_2_ depletion, we purified GIRK2 R92 and its variants (R92Y, R92F, and R92Q) expressed in yeast, and performed a K⁺ flux fluorescence assay with channels reconstituted into liposomes containing POPE and POPG but no PIP_2_^32^ (**Figure 3A,B**). In this assay, application of the proton ionophore carbonyl cyanide m-chlorophenyl hydrazone (CCCP) allows H^+^ to flux into the liposome if GIRK2 channels are open and quenches 9-Amino-6-chloro-2-methoxyacridine (ACMA) fluorescence. To measure the total capacity of K^+^ flux in the liposome, we add the K^+^ ionophore valinomycin at the end of the experiment (**Figure 3B**). We investigated the effect of different concentrations of diC8-PIP_2_ (3, 7, 10, and 50 μM) on activating GIRK2 channels and used the GIRK2 inhibitor phenyl-methanethiosulfonate (MTS-F) to block channel activity^37^. With R92, we observed little fluorescence quenching without PIP_2_ (**Figure 3C**, panel i, black trace), indicating little or no channel activity. Increasing concentrations of PIP_2_ led to dramatic enhancement of quenching, consistent with PIP_2_-dependent channel activation (**Figure 3C**, panel I, teal traces). For R92Y, Q, and F channels, we observed fluorescence quenching even in the absence of PIP_2_ (**Figure 3C**, panels ii-iv, black traces). For R92Y, fluorescent quenching increased with increasing concentrations of diC8-PIP_2_ (**Figure 3C**, panel ii, teal traces), indicating additional PIP_2_ activation. R92Q channels displayed little to no change in quenching with increased PIP_2_ concentrations (**Figure 3C**, panel iii, teal traces). Strikingly, R92F channels exhibited robust quenching upon CCCP addition with no additional changes in the rate of quenching with different concentrations of diC8-PIP_2_ (**Figure 3C**, panel iv), indicating no activation with PIP_2_. To quantify these changes, we constructed dose-response curves using the fluorescence decay rates and plotting as a function of PIP_2_ concentration (**Figure 3D**, panel i). R92Y exhibited a higher EC_50_ for diC8-PIP_2_ than R92 channels (8.4 μM and 25.8 μM, respectively). However, R92Q and R92F appeared insensitive to diC8-PIP_2_, indicated by little change in flux with different concentrations of diC8-PIP_2_ (**Figure 3D**, panel ii). Taken together, all three (R92Y, R92F, and R92Q) substituted channels exhibited intrinsic fluorescence quenching in the absence of any PIP_2_, consistent with what would be expected for a PIP_2_-independent gating mechanism. Furthermore, it is unlikely other phosphoinositides contribute to the channel activity since no other PIs are present in the liposomes (**Figure 3B**).

**Figure 3.**
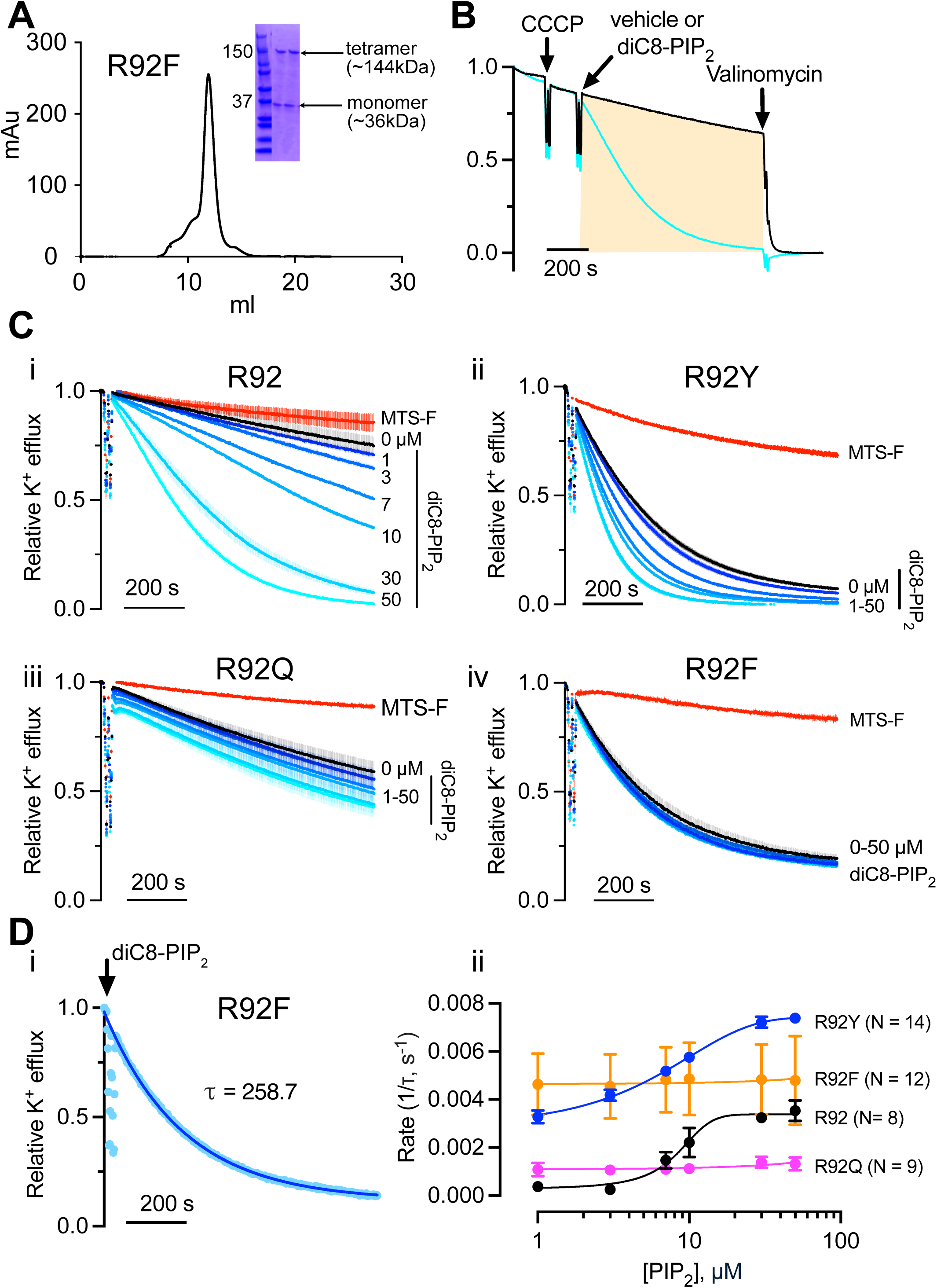
Functional characterization of purified R92 substitutions reconstituted for K^+^ efflux assays. (**A**) Graph shows size-exclusion chromatography trace for purified R92F channels, with peak corresponding to GIRK2 tetramers (∼144 kDa). Inset, SDS-PAGE gel shows the presence of both tetrameric and monomeric (∼36 kDa) channels. (**B**) Representative example of K^+^ efflux assay with purified GIRK2 channels reconstituted into liposomes. Addition of CCCP (protonophore) supports K^+^ efflux, addition of diC8-PIP_2_ accelerates efflux, and valinomycin (K^+^ ionophore) reveals maximal K^+^ efflux for normalization. (**C**) Impact of R92 substitutions on K^+^ efflux and PIP_2_ sensitivity. (i) K^+^ efflux (normalized fluorescence quenching) for GIRK2 is plotted as a function of time showing channel activation in the presence of increasing concentrations of diC8-PIP_2_ (0-50 µM). Basal flux is inhibited with application of MTS-F (100 µM, red trace). N=8. (ii) K^+^ efflux for R92Y (N=14), (iii) R92Q (N = 9), and (iv) R92F (N= 12). (**D**) Analysis of K^+^ efflux rate and diC8-PIP_2_ sensitivity with different R92 substitutions: (ii) Representative example showing rate of flux for GIRK2 derived from an exponential fit to determine the tau. (ii) The rate of K^+^ efflux (1/tau) plotted as a function of diC8-PIP_2_ concentration for R92 and substitutions. Lines show best fit using the Hill equation with EC_50_ of 29 µM and 13 µM for R92 and R92Y, respectively. Lines show best fit using the simple linear regression for R92F and R92Q. Mean ± SEM plotted with indicated N.

### Molecular dynamics simulations suggest that PIP_2_ interaction is weakened in R92F

The combination of results from patch-clamp electrophysiology and K^+^ flux assays with reconstituted GIRK2 channels provide evidence that these substitutions at R92 alter the interaction of PIP_2_ with the channel. To further investigate this possibility, we conducted molecular dynamics (MD) simulations using models of GIRK2 R92 and R92F (**Figure 3D**, panel ii). We docked PIP_2_ in the PIP_2_ binding pocket based on the GIRK2 structure (PDB: 4KFM) and conducted simulations for 400 ns. We also conducted three independent simulations (**Supplementary Figure 1**). Compared to R92 simulations, the root-mean-square fluctuation (RMSF) increased in the slide helix (residues 66-92) and TM1 (residues 92-100) of R92F, suggesting greater flexibility in these regions in the R92F channel (**Figure 4A**, green highlights). This enhanced flexibility suggests the slide helix adopts a more relaxed conformation compared to R92 channels. Moreover, the TM1 domain moves away from TM2 toward the end of the 400 ns simulation, coinciding with the dissociation of PIP_2_ from its binding site (**Figure 4A**, blue highlight).

**Figure 4.**
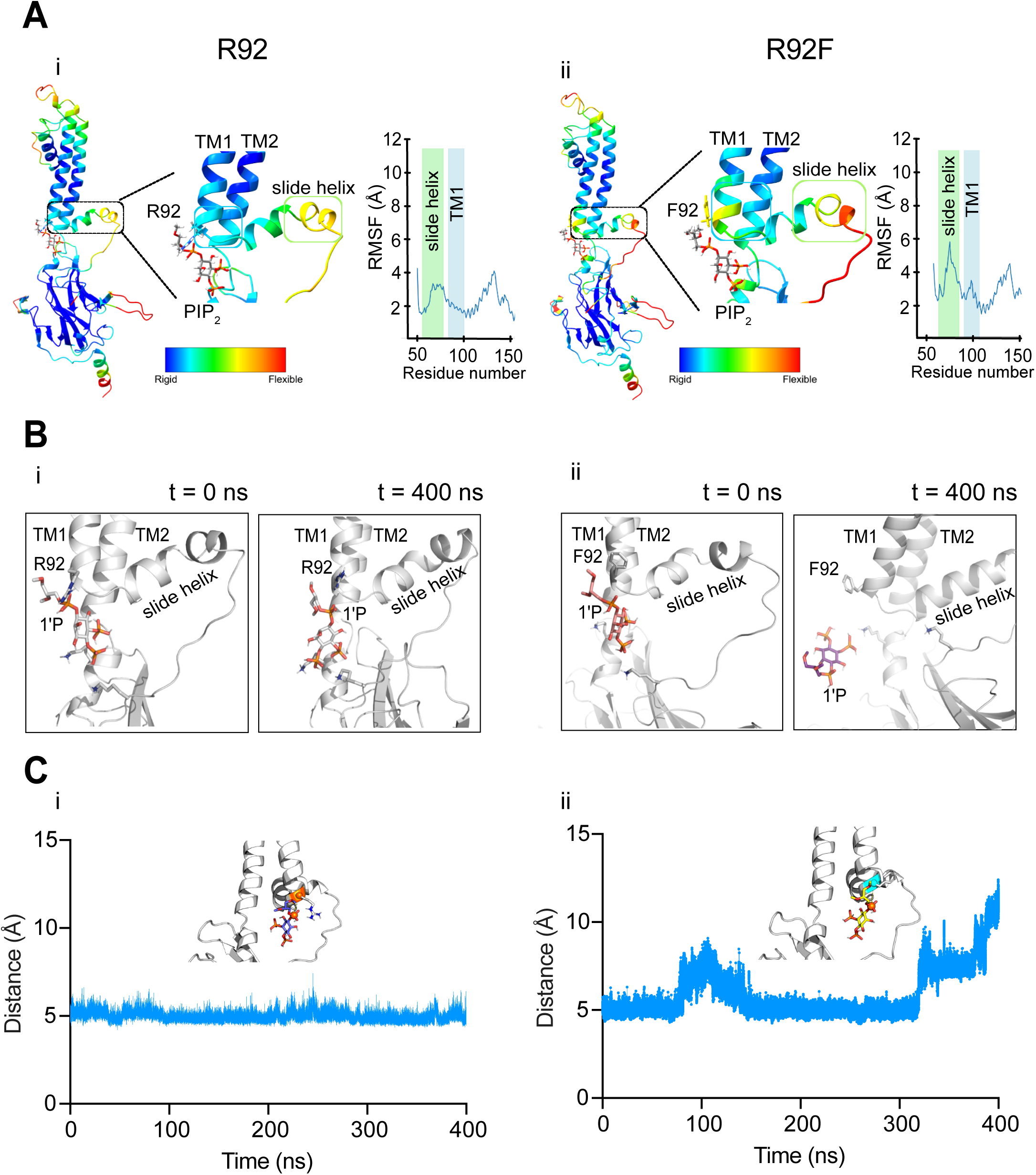
Molecular dynamics simulation of R92 and R92F channels with PIP₂. (**A**) Structures of R92 (i) and R92F (ii) single subunit from MD1 show backbone flexibility, color-coded for root mean square fluctuation (RMSF) (rigid regions in blue, flexible in red). Boxed regions show expanded views of R92 or R92F with PIP₂. Plots show RMSF highlighting the slide-helix and TM1 transmembrane domain for both R92 and R92F. (**B**) Snapshots of MD1 for R92 (i) and R92F (ii) with PIP₂ at initial (t = 0 ns) and final (t = 400 ns) time points of the simulation. (**C**) Time-resolved analysis of the distance between the C-alpha carbon of R92 (i) or R92F (ii) and the 1’ PO_4_ of PIP₂ over MD1. Inset shows snapshots of R92/PIP₂ or R92F/PIP₂ interactions, respectively.

A detailed analysis of the PIP_2_ binding site revealed that the interaction of R92F with the 1′-phosphate group (1′-PO₄) of PIP_2_ is weakened, destabilizing PIP_2_ within the PIP_2_ binding pocket of GIRK2 (Figure 4B, panel i, ii). At the start of the MD simulation, the 1′-PO₄ group of PIP_2_ forms hydrogen bonds with the backbone of R92F (**Figure 4B**, panel ii, t = 0 ns). As the simulation progresses, the hydrophobic side chain of R92F disrupts interactions with the 1’-phosphate group, releasing PIP_2_ from the R92F structure (**Figure 4B**, panel ii, t = 400 ns). We also measured the distance between the Cα in R92 and R92F and the 1’-phospho atom of PIP_2_ over the simulation time. For R92, the distance between R92 and 1′-PO₄ of PIP_2_ remained the same during the simulation (**Figure 4C**, panel i) while the distance between R92F and the 1′-PO₄ increased after ∼ 300 ns of the MD simulation (**Figure 4C**, panel ii). Compared to the R92, R92F also formed fewer hydrogen bonds with PIP_2_, with the hydrophobic side chain appearing to destabilize the PIP_2_ in the pocket as the simulation progressed (**Supplementary Figure 2**). These observations are consistent with R92F channels exhibiting an insensitivity to PIP_2_ in the functional assays. Together, these findings provide a plausible hypothesis for the PIP_2_-independent phenotype observed in the R92 substitutions, whereby the slide helix is more flexible and allow PIP_2_ to diffuse out of the PIP_2_ pocket.

### Structural Characterization of R92F in GIRK2 Channels

If PIP_2_ is destabilized in the PIP_2_ pocket of GIRK2, as the MD simulations suggest, then how does the channel open in the absence of PIP_2_? One possibility is that R92F somehow mimics a PIP_2_-bound state. To explore this further, we used single-particle cryo-EM to determine the structures of R92F and R92Q in the absence of PIP_2_. Analysis of R92F particles revealed two distinct conformational states. State 1, resolved at 3.68 Å, reveals a more tilted CTD relative to the TMD (**Figure 5A**, panel i). State 2 has a global resolution of 3.62 Å, showing an overall less tilted CTD (**Figure 5A**, panel ii). Both reconstructions were solved with C1 symmetry due to the inherent asymmetry in R92F structures. Similarly, we analyzed R92Q channels and observed two particle populations, categorized into conformational states resembling those seen in R92F (**Supplemental Figure 4**). The bottom views of all four maps suggest a constricted G-loop gate, suggesting the structures of both R92F and R92Q channels in detergent micelles represent closed conformations.

**Figure 5.**
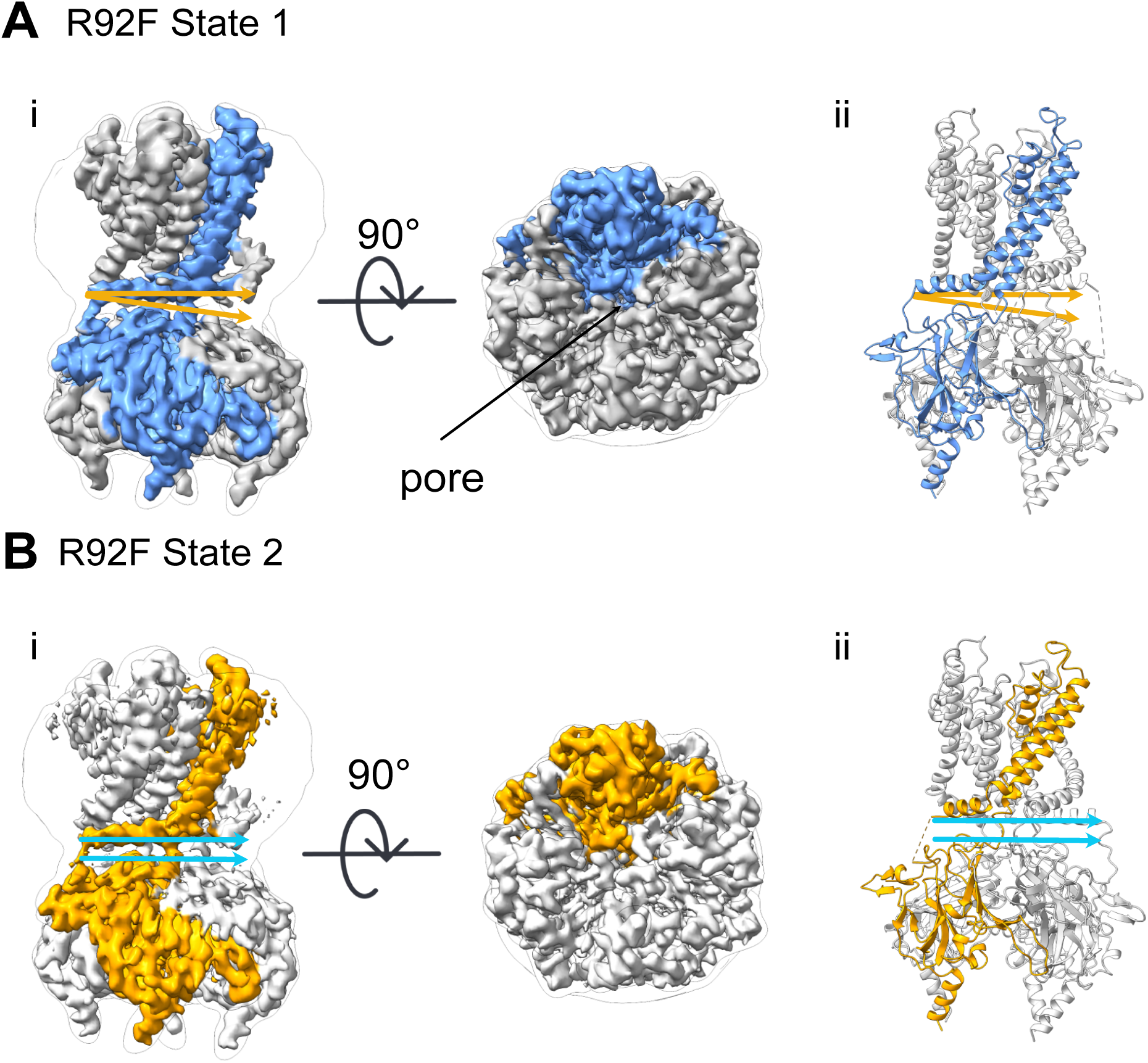
Cryo-EM structures of GIRK2 R92F channels in two conformational states. **(A)**(i) Cryo-EM map of R92F State 1, with a global resolution of 3.68 Å, shown from two orientations: side view (left) and bottom-up view (right). (ii) Atomic model of R92F State 1 built from the reference model 6XEU. (**B**) (i) Cryo-EM maps of R92F State 2, with a global resolution of 3.62 Å, shown from two orientations: side view (left) and bottom-up view (right). (ii) Atomic model of R92F State 2 built from the model of R92F State 1. The tilt levels of the CTD relative to the slide helix in State 1 and State 2 are demonstrated by arrows.

From the 3D maps, we built structural models for both states of R92F (**Supplementary Table 1**). For our structural comparisons, we focused on state 1 due to its higher resolution of the slide helix. We aligned state 1 of R92F (PDB:9MH9) with the apo GIRK2 structure (PDB: 6XIS) and the PIP_2_-bound GIRK2 structure (PDB: 6XEU) (**Figure 6A**). R92F induces significant conformational shifts compared to the native channel: on one side of the tetramer, the CTD-TMD distance is 17 Å (measured between the C-alpha atoms of L229 and T80), resembling the CTD-TMD proximity in the PIP_2_-bound state (referred to as the CTD-engaged or docked conformation). In the other side of the tetramer, the CTD-TMD distance is 24 Å, similar to the apo state, representing a CTD-detached or undocked conformation (**Figure 6A**). These findings suggest that the R92F substitution stabilizes an intermediate state between the apo and PIP_2_-bound conformations, where the CTD is only partially engaged with the TMD.

**Figure 6.**
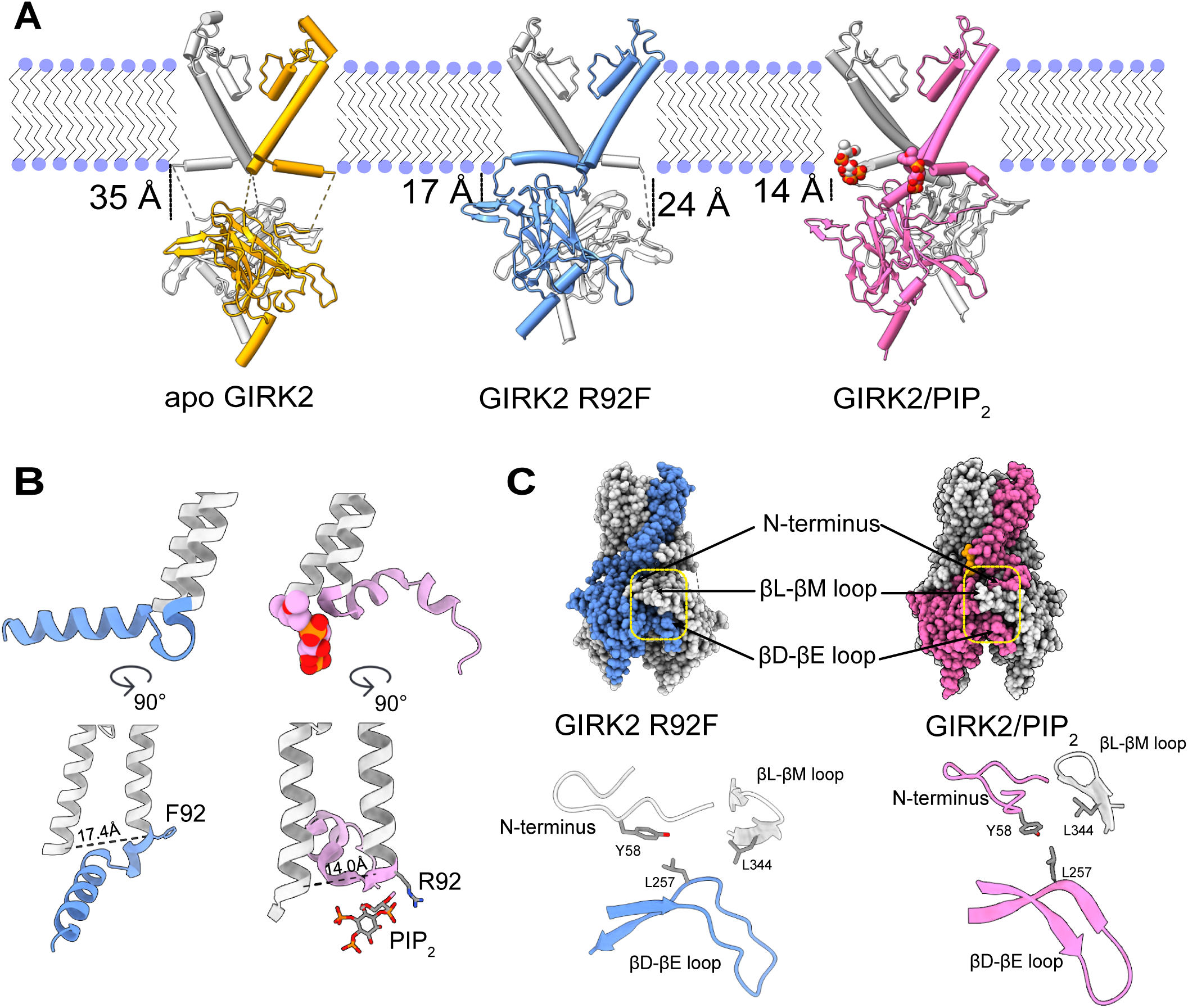
Structural comparison of R92F and R92 complexes with and without PIP_2_. (**A**) Side view of two representative subunits for GIRK2 in the apo state (yellow, left, PDB:6XIS), R92F apo (blue, middle, PDB:9MH9), and GIRK2 with PIP2 (pink, right, PDB:6XEU). The distances between the transmembrane domains and the CTD are shown, with 35 Å for apo GIRK2, 17 Å and 24 Å for R92F, and 14 Å for GIRK2/PIP_2_. (**B**) Comparison of slide-helix in R92F (blue, left) and GIRK2/PIP_2_ (pink, right). The distance between the transmembrane helices (T80 on TM1 and I195 on TM2) is shown (17 Å for R92F and 14 Å for GIRK2/PIP_2_), along with the specific interaction of the R92 residue with the 1’-PO_4_ of PIP_2_. (**C**) Top: Surface representations of R92F (blue, left) and GIRK2/PIP_2_ complex (pink, right) with annotations for the alcohol/Gβγ interaction site (the N-terminus, βL-βM loop, and βD-βE loop regions). Bottom: Zoom of the alcohol/Gβγ interaction site, highlighting the interactions between Y58 (alcohol), L257 (alcohol), and L344 (Gβγ) in two adjacent subunits.

The C1 asymmetry observed may indicate the CTD undergoes a continuous rocking motion of that is not fully captured by discrete classes in cryo-EM. It remains possible that R92F may transiently adopt a rare but fully CTD-engaged state. These data suggest that the R92F substitution may mimic the role of PIP_2_ by partially engaging the CTD with the TMD.

### Conformational Changes Induced by R92F Substitution

Comparing the R92F structure to the PIP_2_-bound R92 GIRK2 reveals substantial conformational changes resulting from R92F. Notably, the slide helix (residues 66-92) in the R92F structure is oriented in the opposite direction at Phe-92 compared to Arg-92 in GIRK2 structure (**Figure 6B**). Additionally, the slide helix in R92F (**Figure 6B**, blue) appears straighter and potentially more thermodynamically stable compared to that in GIRK2 R92 (**Figure 6B**, pink). The R92F generates an electron cloud that repels the electron density of adjacent residues (W91 and K90), reorienting the slide helix such that it protrudes into the region where PIP_2_ normally binds, potentially preventing PIP_2_ from interacting with the channel (**Figure 6B**). This may explain why R92F channels reconstituted into liposomes are unresponsive to diC8-PIP_2_ in K^+^ flux assays.

Moreover, this reorientation of the slide helix occurs concomitantly with a bending of TM2 moving it away from TM1, as indicated by the increased distance between T80 (on TM1) and I195 (on TM2), which expands from ∼14 Å in the wild-type (Figure 6B, pink) to ∼17 Å in the R92F structure (**Figure 6B**, blue). This increased TM1-TM2 distance aligns with the enhanced flexibility of TM1 observed in MD simulations (**Figure 4A**, panel ii). The widening of the TM1-TM2 distance also constricts the inner helix gate of the R92F channel compared to the wild-type, suggesting the R92F channel is captured in a closed conformation.

The reorientation of the slide helix in the R92F structure also alters the inter-subunit interaction profile. In GIRK2 R92, the N-terminus (residues 55-70) interacts closely with the βD-βE loop of the same subunit and the βL-βM loop of an adjacent subunit (**Figure 6C**, pink), forming a hydrophobic pocket where alcohol is likely activates the channel^3,36^. However, in R92F, the N-terminus and the βL-βM loop are derived from the same subunit while the βD-βE loop is contributed from the adjacent subunit, leading to a novel interaction between the N-terminus and the βD-βE loop (**Figure 6C**, blue). This rearrangement alters the native alcohol-binding pocket, which is now composed of different subunit regions. Additionally, the βL-βM loop, where Gβγ subunits bind^39^, is displaced in the R92F structure, potentially impairing Gβγ-mediated interactions with the channel (**Figure 6C**, blue). These conformational changes could explain the altered responses of R92F channels to activation with propanol and baclofen observed in electrophysiological experiments in HEK cells (**Figure 1C**, panel ii).

In conclusion, the structural analysis of R92F and R92Q highlight the critical role of arginine at position 92 in maintaining the correct orientation of the slide helix and inter-subunit interactions for proper gating by PIP_2_, alcohol, and G proteins. The R92F structure induces conformational changes in the TMD-CTD interface that partially mimic the PIP_2_-bound state, offering interesting insights into the structural dynamics governing GIRK2 channel regulation.

## Discussion

### A putative PIP_2_ Independent Gating of GIRK2 Channel

Prior studies demonstrated that PIP_2_ is crucial and indispensable for Kir channel activation. For example, the interaction of PIP_2_ with Kir6.2 reduces the ATP sensitivity of the channel, thereby affecting its ATP-dependent closure^40^. Another example is that PIP_2_ plays a role in the pH inhibition of Kir4.1/Kir5.1 heterotetramers^41,42^. Lastly, a recent cryo-EM structural study of Kir4.1 homotetramers revealed that PIP_2_ binds to a similar pocket as in GIRK2, with the 1’-phosphate of PIP_2_ interacting with the R92 equivalent in Kir4.1, R65^43^. A mutation of Kir4.1 (R65P) is found in SeSAME/EAST syndrome, a channelopathy associated with seizures, deafness, ataxia, and developmental abnormalities^44^. R92P results in non-specific binding across various PIP variants^34^. These studies underscore the essential role of PIP_2_ in regulating different Kir subtypes. However, our results with GIRK2 R92Y, R92F, and R92Q channels provide the first evidence of functional GIRK channels operating in the absence of PIP_2_. These substitutions, located at the PIP_2_ binding site, alter the regulation of GIRK channels by this membrane lipid. Although the exact mechanism by which these GIRK2 mutants function independently of PIP_2_ is unknown, several hypotheses can explain this unique property.

One possibility is that the chemical characteristics of the substituted side-chains; the aromaticity of tyrosine and phenylalanine, the polarity of tyrosine and glutamine and the lack of charge on all substitutions-allow these channels to interact with other membrane phospholipids besides PIP_2_, promoting K^+^ conduction. A lipid mechanism has been proposed for substitutions at the K62 in Kir2.2where a nonpolar, aromatic amino acid (W) substitution at the membrane interface interacts with bulky anionic phospholipids to enhance PIP_2_ sensitivity and reduce the gap between the CTD and TMD^45^. In GIRK2, the bulky amino acid substitutions at R92 may facilitate a similar process with lipids, drawing the CTD closer to the membrane and even directly opening the channel. Future studies with GIRK2 reconstituted in nanodiscs containing different phospholipids could help characterize further this possible mechanism of PIP_2_-independent activity.

Another possible explanation is that the cryoEM structures of GIRK2 R92F and R92Q reveal a constricted HBC gate in the absence of PIP_2_, but are open sufficient for K^+^ to traverse the channel in their partially dehydrated form. An MD simulation study on Kir channels suggested that unhydrated K^+^ passing through the selectivity filter of Kir channels are rehydrated with 6-7 water molecules in the inner vestibule^46^. When approaching the HBC gate, these hydrated K^+^ are then partially dehydrated, allowing passage through a constricted gate, even in the absence of PIP_2_^46^.

### A refined Model for PIP_2_ Gating in GIRK Channels

In the absence of PIP_2_, the CTD becomes disengaged from the TMD in the GIRK2 structure^29,31^. The slide helix is poorly resolved, however, indicating significant conformational flexibility in this region^29,31^. Furthermore, this region is usually truncated due to its disordered nature to improve the expression in a heterologous system^32^. Thus, the poor resolution makes it difficult to determine whether the slide helix only adopts one orientation in both the apo and PIP_2_-bound structures. A potential gating mechanism could involve the slide helix switching orientations dynamically in the closed state, contributing to its lower resolution compared to the rest of the protein. In native channels, PIP_2_ stabilizes the slide helix in an orientation conducive to PIP_2_ binding (right-sided slide helix). In contrast, the cryo-EM structures of GIRK2 R92F and R92Q show that the slide helix appears in a conformation that may prevent PIP_2_ from binding to the channel (left-sided slide helix). In both cases, the CTD moves closer to the TMD and the membrane interface, priming the channel for opening and K^+^ conduction (**Figure 7**).

**Figure 7.**
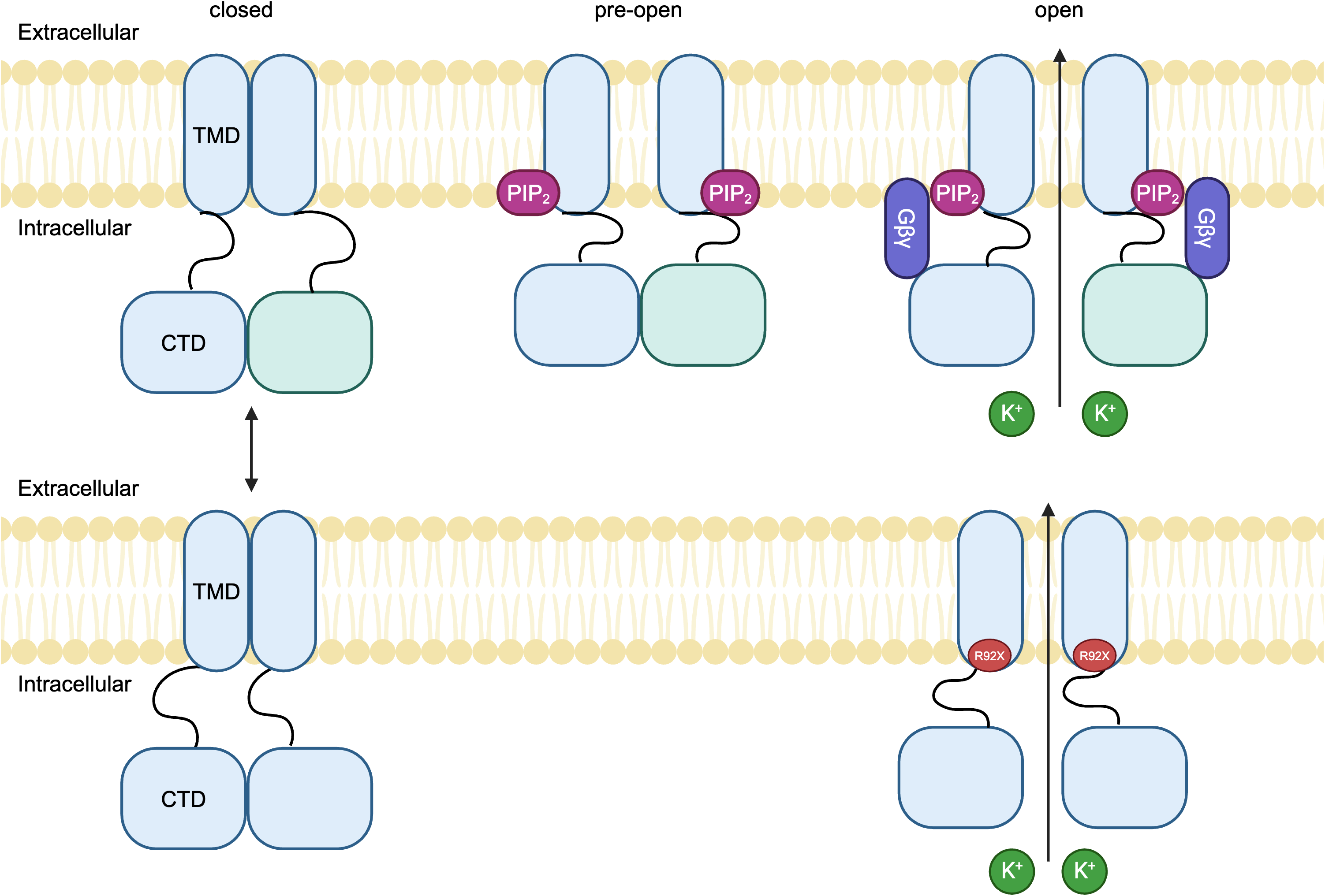
Proposed model for the role of R92 in GIRK2 channel activation. Top: GIRK2 channels adopt the right-facing slide helix (R). In the absence of PIP_2_ and Gβγ, GIRK2 channels remain closed, with the transmembrane domain (TMD) and cytoplasmic domain (CTD) loosely associated. Binding of PIP_2_ to the channel promotes a pre-channel opening. Subsequent binding of Gβγ stabilizes the open state, allowing enhanced K^+^ efflux. Bottom: GIRK2 channels adopt the left-facing slide helix (L). In the case of R92Y, R92Q, and R92F, the slide helix is locked in this conformation. The R92X substitutions are proposed to open the channel in a PIP_2_-independent manner.

Single particle analyses of GIRK2/PIP_2_ complexes reveal a dynamic range of CTD-TMD engagement, with some particles adopting a tilted CTD, suggesting that PIP_2_ binding does not ensure CTD-TMD engagement^29,31^. Our structures of R92F and R92Q without PIP_2_ exhibited a similar tilted CTD conformation seen in a subset of GIRK2/PIP_2_ particles. This observation is corroborated by a cryo-EM structure of human Kir2.1 channels, which demonstrates that CTD-TMD tethering can occur independently of PIP_2_^47^. Collectively, these findings suggest a refined gating mechanism in GIRK2, where the CTD undergoes a “rocking movement” in the absence of PIP_2_, intermittently engaging with the TMD. The primary function of PIP_2_ may be to stabilize the tethering between the two domains, rather than initiating the engagement of the CTD with the TMD.

### Physiological Significance of R92 in GIRK2 Channel Gating

Our results show that R92 plays a critical role in the binding of PIP_2_ in GIRK2. In addition, our results underscore the importance of R92 in endowing GIRK channels with the property of gating by both Gβγ and alcohol; R92 ensures a specific arrangement of the subunit interface in the CTD, with the βD-βE loop and N-terminus from one subunit, and the βL-βM loop from the neighboring subunit creating the Gβγ and alcohol pocket. Interestingly, R92 is conserved across most Kir channels, except for Kir6.2, which has a proline^48^. Substitutions in this region in Kir channels have been linked to rare diseases such as SeSAME/EAST syndrome caused by mutations in Kir4.1 channels^44^. One disease-associated mutation, R65P, decreases the open probability and pH sensitivity of Kir4.1 channels and is homologous to GIRK2 R92^44,49^. Interestingly, R92P makes GIRK2 more promiscuous to gating by different PIP isoforms^34^. Taken together, these studies suggest that R92 and the structural properties of its side chain are critical for binding PIP_2_ and maintaining channel activity. The significance of R92 may extend to other Kir channels, where equivalent residues have been implicated in mediating the interaction with PIP_2_ and regulating channel opening^47,50^. Perhaps substitutions at this equivalent position in other Kir channels also confer PIP_2_-independent gating.

In addition to its role in coordinating PIP_2_-mediated gating, R92 is a key site for cholesterol regulation of GIRK channels^29^. The binding of CHS in the presence of PIP_2_ increases the proportion of CTD-TMD-engaged particles in single-particle analyses, highlighting the importance of R92 in cholesterol-mediated potentiation of GIRK channels^29^. Even though cholesterol sites have not been structurally characterized in other Kir channels, given the structural conservation of this family, it would be interesting to examine the role of the amino acid homologous to R92 in cholesterol-mediated gating of other Kir channels.

In summary, R92 plays a crucial structural role in regulating PIP_2_ binding and maintaining proper channel function for gating by Gβγ and alcohol. This residue’s interactions with both PIP_2_ and cholesterol are essential for coordinating the gating mechanism, highlighting its significance in channel modulation. Understanding the molecular mechanisms governing R92 interactions in GIRK2 provides critical insights that may help enable the development of targeted GIRK channel therapeutics^25^.

## Methods

### Molecular Biology and Cell Culture

Single point mutations at R92 (A, W, F, Y, K, Q) were introduced into mouse GIRK2c using site-directed mutagenesis (QuickChange II XL, Agilent Technology) and confirmed by DNA sequencing. Human Embryonic Kidney 293T (HEK293T) cells were maintained in Dulbecco’s modified Eagle’s medium (DMEM; Sigma-Aldrich, St. Louis, MO, USA) supplemented with 10% (v/v) Fetal Bovine Serum (FBS), 100 U/ml penicillin, 100 mg/ml streptomycin, and 1X Glutamax (ThermoFisher) in a humidified 37°C incubator with 5% CO_2_. Cells were plated in 12-well plates and transiently transfected with cDNA (GIRK2c WT or mutants: 0.5 μg; eYFP: 0.02 μg, for identification of transfected cells; GABA_B1b_: 0.2 μg; GABA_B2_: 0.2 μg; Dr-VSP: 0.4 μg) using Lipofectamine 2000 (ThermoFisher). After 48 hours, cells were transferred onto poly-D-lysine-coated (100 mg/ml; Sigma-Aldrich) 12-mm glass cleaned coverslips in 24-well plates for whole-cell patch-clamp recording^29^.

### Electrophysiology

HEK293T cells were transiently transfected with GIRK2 or the R92 substitution cDNA, and whole-cell patch-clamp recordings were performed as previously described^36^. Briefly, borosilicate glass electrodes (Warner Instruments) with a resistance of 3–6 MΩ were used, filled with an intracellular solution composed of 130 mM KCl, 20 mM NaCl, 5 mM EGTA, 5.46 mM MgCl_2_, 2.56 mM K_2_ATP, 0.3 mM Li_2_GTP, and 10 mM HEPES (pH 7.4, ∼320 mOsm). The extracellular ‘20K’ solution contained 20 mM KCl, 140 mM NaCl, 0.5 mM CaCl_2_, 2 mM MgCl_2_, and 10 mM HEPES (pH 7.4, ∼310 mOsm). Currents were evoked at 0.5 Hz using a voltage step to −120 mV from a holding potential of −40 mV, followed by a ramp voltage protocol (−120 mV to +50 mV, E_K_ = −50 mV with 20 mM K_out_). K^+^ currents were corrected for series resistance and measured at −120 mV with either the 20K solution or the 20K + Ba^2+^ (1 mM) solution. The basal current was defined as the Ba^2+^-sensitive current.

To study the effect of membrane PIP_2_ depletion, Dr-Vsp was activated by applying a voltage step from a holding potential of −40 mV to +100 mV for varying durations, followed by measuring the amplitude of GIRK currents at −120 mV. This three-step protocol was repeated every 2 seconds^37^. The degree of Dr-Vsp inhibition was calculated as a fraction of the Ba^2+^-sensitive basal current. All data are presented as mean ± S.E.M, and statistical significance (P < 0.05) was evaluated using one-way ANOVA.

### Protein purification and liposome reconstitution

All GIRK channels were expressed and purified in *Pichia pastoris* as previously described^27^. In brief, the highest-expressing clone was cultured in BMGY medium and induced in BMM medium containing 1% methanol. Cells were harvested, resuspended in buffer (50 mM HEPES, pH 7.4; 150 mM KCl; 1 mM TCEP; 1 mM AEBSF and Complete EDTA-free protease inhibitor tablets (Roche)), flash frozen in liquid nitrogen, and stored at −80°C. Frozen cells were lysed using a Mixer Mill (Retsch) with five cycles of 3 minutes at 25 Hz and stored as powder at −80°C until use. The cell powder was then solubilized in buffer containing 50 mM HEPES, pH 7.35; 150 mM KCl; 1 mM TCEP; 1 mM AEBSF; Complete ULTRA EDTA-free protease inhibitor tablets (Roche) and 2% (w/v) n-Dodecyl-β-D-maltoside (DDM; Anatrace) with gentle stirring at 4°C. Unsolubilized material was removed by centrifugation at 40,000 × g for 40 minutes at 4°C and then filtered. The supernatant was incubated with HISPur Cobalt charged resin (ThermoFisher) equilibrated in wash buffer (50 mM HEPES, pH 7.0; 150 mM KCl; 0.2% DDM; 20 mM imidazole). The resin was then washed with 10 column volumes (CV) of wash buffer, followed by 5 CV containing 40 mM imidazole, and eluted in a buffer containing 300 mM imidazole. The eluate was pooled, exchanged into an imidazole-free buffer, and digested overnight at 4°C with HRV 3C protease^51^. The protein was subsequently concentrated and subjected to size exclusion chromatography on a Superdex-200 gel filtration column in buffer containing 20 mM TRIS-HCl pH 7.5, 150 mM KCl, 20 mM DTT, 3 mM TCEP, 1 mM EDTA, and 0.025% (w/v) DDM (anagrade). Fractions eluting at a volume corresponding to the GIRK channel tetramer were pooled, concentrated, and analyzed by SDS-PAGE and Coomassie blue staining.

The purified GIRK2 channels were reconstituted into lipid vesicles as described previously^27^. In brief, a lipid mixture consisting of 1-palmitoyl-2-oleoyl-sn-glycero-3-phosphoethanolamine (POPE), 1-palmitoyl-2-oleoyl-sn-glycero-3-phospho(1’-rac-glycerol) (POPG), at mass ratios of 3:1 was prepared, reconstituted in vesicle buffer (20 mM K-HEPES, pH 7.4; 150 mM KCl; 0.5 mM EDTA containing 35 mM CHAPS), and incubated with protein in detergent at a 1:200 protein ratio. Detergent removal was achieved through the sequential addition of Bio-beads SM-2 (Bio-Rad). All phospholipids were obtained from Avanti Polar Lipids, Inc. Water-soluble PIP_2_ (diC8-PIP_2_) was purchased from Echelon Biosciences.

### Flux assay

The fluorescence-based flux assay for GIRK channel activity was conducted as previously described^27,32^. In brief, liposomes were diluted 1:20 into a flux buffer (20 mM Na-HEPES, pH 7.4; 150 mM NaCl; 0.5 mM EDTA) containing 5 μM of the H^+^ sensitive dye 9-Amino-6-chloro-2-methoxyacridine (ACMA) (Invitrogen). Fluorescence measurements were taken using a Flexstation 3 microplate reader (Molecular Devices) with the following settings: 410 nm excitation, 480 nm emission, 455 nm cutoff, medium PMT sensitivity, and 2-second sampling intervals. After a stable baseline fluorescence was established (150 s), the H^+^ ionophore m-chlorophenyl hydrazone (CCCP) (Sigma) was added automatically (1 μM final concentration), followed by addition of either vehicle or benzyl methanethiosulfonate (MTS-F, 100 μM final; Toronto Research Chemicals) 150 seconds later, and lastly the addition of the K^+^ ionophore Valinomycin (100 nM final; Invitrogen) 900 seconds after that, to determine the maximum K^+^ flux. GIRK2 channels are likely oriented in both directions within the liposomes; however, channels oriented inside-out are expected to support high K^+^ flux due to the high Na^+^ in the flux buffer and high K^+^ inside the liposome^27^. The percentage of relative K^+^ flux (or relative fluorescence intensity) was calculated by measuring the extent of quenching 10 seconds before adding Valinomycin and dividing it by the total quenching capacity of the liposomes, normalized to the relative fluorescence units (RFU) 10 seconds before the addition of vehicle or MTS-F (F0). The liposome flux assay illustration was created using Biorender.com.

### Data Analysis

Fluorescence measurements were normalized to compare different flux assay experiments, using either the baseline fluorescence prior to CCCP (F_b_) and the fluorescence after Valinomycin (F_v_), or in the case of acute compound addition, the fluorescence post-CCCP (F_b_) and after Valinomycin (F_v_). The normalized fluorescence (F_N_), referred to as “relative K^+^ flux,” was calculated using the following equation (Eq. 1):

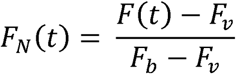

where F_N_(t) represents the normalized fluorescence at time t, and F(t) is the fluorescence intensity as a function of time.

The rate of K^+^ flux (1/τ) was determined by fitting the decay of normalized fluorescence with a single exponential equation (Eq. 2):

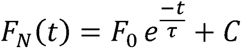

where F_0_ is the amplitude, 1/τ is the flux rate, and C is a constant. In cases where fitting the decay was not feasible, the amplitude of fluorescence just before adding Valinomycin (1190 s) was used to determine the fractional inhibition of K^+^ flux.

Dose-response curves for diC8-PIP_2_ were fitted using the Hill equation (Eq. 3):

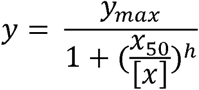

where y_max_ is the maximum flux rate or fractional inhibition, [X] represents the concentration, h is the Hill coefficient, and x_50_ is the half-maximal effective concentration (EC_50_ for PIP_2_). All data are presented as mean ± S.E.M, and statistical significance (P < 0.05) was evaluated using one-way ANOVA followed by Dunnett’s multiple comparison test.

### Molecular dynamics (MD) simulation

The PDB: 4KFM crystal structure of the GIRK2 channel bound to a PIP_2_ head group served as the basis for all MD simulations, following previous work^33^. The GIRK2-PIP_2_ complex was embedded into a bi-layer model using CHARMM-GUI^52^. The R92 system included 424 POPC lipids and 0.15M KCl ions to achieve charge neutrality within a triclinic periodic box measuring 131×131×102 nm. KCl ions were added to achieve charge neutrality and a salt concentration of 0.15 M. The CHARMM36m force field was applied to the protein, lipids, and ions, while water molecules were modeled using the TIP3P model^53^. A second system (R92F) was created by generating the R92F mutation via CHARMM-GUI in silico mutagenesis.

All MD simulations were conducted using the GROMACS simulation package version 2020.3^54^. Following the standard CHARMM-GUI protocol, each system underwent a 100-nanosecond equilibration phase, which included multiple consecutive simulations in the NVT and NPT ensembles with harmonic restraints gradually removed from the lipids, ligand, and protein atoms. Next, unrestrained production MD simulations were carried out for 400 ns for each system, with simulation coordinates saved every 100 ps. The system’s temperature was maintained at 310K using the Nosé-Hoover scheme, and the pressure was kept constant at 1 atm using a Parrinello-Rahman approach with semi-isotropic scaling. This equilibration production and simulation was repeated for both the R92 and R92F systems using independent replicas.

### Analysis of simulation trajectories

The simulation trajectories were performed using MDAnalysis^55^. The root mean square deviation (RMSD) and root mean square fluctuation (RMSF) were calculated using the fast QCP algorithm^55^. Distances between the α-carbon of R92F and the phosphorus atom of 1’PO_4_ of PIP_2_, as well as the α-carbon of R92 and the phosphorus atom of 1’PO_4_ of PIP_2_, were measured for the length of the simulation time. The g_hbond function in GROMACS was utilized for hydrogen bond analysis between R92F and PIP_2_ as well as R92 and PIP_2_, with the default settings of a 3.5 Å cutoff distance and a 30° angle.

### CryoEM sample preparation, data collection, and processing

The quality of purified samples was initially assessed using negative stain EM according to established protocols^56^. For cryoEM analysis, 3 μl of GIRK2 mutant (R92F and R92Q) at a concentration of 7–10 mg/ml were applied to glow-discharged Quantifoil Au1.2/1.3, 200 mesh grids. The grids were then blotted and plunge-frozen in liquid ethane using an FEI Vitrobot. R92Q cryoEM data were collected at 300 kV on a Titan Krios microscope equipped with a Gatan K3 direct detection camera. Raw images were captured as movies, recorded at a magnification of 81000x, corresponding to a pixel size of 1.058 Å at the specimen level. Each movie was recorded over 2 seconds with a frame rate of 0.05 seconds per frame, resulting in a total dose of 51.73 electrons per pixel, with defocus values ranging from −0.1 to −3.1 μm. R92F cryoEM data were collected at 300 kV on a Titan Krios microscope equipped with a Gatan K3 direct detection camera at the New York Structural Biology Center. Raw images were captured as movies, recorded at a magnification of 81000x, corresponding to a pixel size of 1.069 Å at the specimen level. Each movie was recorded over 2 seconds with a frame rate of 0.05 seconds per frame, resulting in a total dose of 53.92 electrons per pixel, with defocus values ranging from −0.8 to −2.2 μm.

The micrographs were motion-corrected and dose-weighted using Patch Motion Correction in cryoSPARC. Defocus estimation was performed using Patch CTF Estimation. Particles were picked using a blob picker with a diameter of 100–150 Å, resulting particles were subjected to two rounds of 2D classification. Initial 3D reconstruction was performed through ab initio reconstruction (k=3), followed by heterogeneous refinement. Selected classes were re-centered and re-extracted for further refinements. Nonuniform refinement with C1 symmetry was applied, including global and local CTF refinement. Subsequent local refinement steps led to high-quality maps. The global resolutions for R92F reconstructions are 3.68 Å (State 1) and 3.62 Å (State 2) and for R92Q reconstructions are 3.78 Å (State 1) and 4.13 Å (State 2). The maps were sharpened with a B-factor of −120 or −140, and local resolution estimates were calculated. The resolutions of the maps were determined using the 0.143 “gold-standard” Fourier Shell Correlation (FSC) criterion. The maps were further processed using DeepEMenhancer to improve interpretability.

### Cryo-EM Modeling and Refinement

The GIRK2/PIP_2_ structure (PDB: 6XEU) was used as an initial model to dock into the cryo-electron microscopy (cryo-EM) density maps of R92F in State 1 and State 2. The density maps were processed and the model was fit using a multi-step approach. First, the 6XEU structure was rigid-body fitted into the cryo-EM density maps of both states using Coot. The secondary structure elements were manually adjusted for optimal fit to the density using this initial rigid-body docking. Following this, further refinement and relaxation of the model into the density was performed with ISOLDE to improve local fitting while maintaining secondary structure restraints to prevent distortion of helices and beta sheets. Where the density map did not provide clear side chain information, side chains were trimmed accordingly to avoid overfitting. The trimming ensured that the model accurately reflected the experimentally observed densities. Final refinement of the models was carried out using PHENIX real-space refinement. Density cutouts from the cryo-EM map were used to focus refinement in areas with high resolution. Parameters included optimizing the placement of backbone and side-chain atoms to best fit the density, with adjustments made to optimize stereochemistry and preserve proper bond lengths and angles. Throughout the entire process, iterative steps of model fitting, refinement, and validation were conducted to ensure the models accurately reflected the observed densities while maintaining structural integrity.

## Data availability

All data needed to evaluate the conclusions in the paper are presented in the paper or the Supplementary Materials. The cryo-EM maps have been deposited to Electron Microscopy Data Bank (EMDB) with accession codes EMD-48268 (R92F State 1), EMD-48267 (R92F State 2), EMD-48270 (R92Q State 1), and EMD-48269 (R92Q State 2). Atomic coordinates have been deposited to the Protein Data Bank (PDB) with accession codes 9MH9 (R92F State 1) and 9MH8 (R92F State 2). The MD trajectory files are freely available at Zenedo.org (https://zenodo.org/) and can be accessed at 10.5281/zenodo.14544811.

## Acknowledgement

We thank members of the Wacker Lab for the cryo-EM discussion. This work was supported by NIH grant 5R01AA018734 and NCCAT-BAG-PS220401

## Contributions

H.N. performed biochemical preparations. H.N., Y.Z. conducted electrophysiology experiments. H.N. and I.G. performed K^+^ fluorescent flux assay. H.N. collected cryo-EM data. J.M. analyzed cryo-EM data. K.H. and Y.Z. ran MD simulations and analysis. P.S. and A.S. conceived and supervised the work. H.N, I.G., and P.S. prepared the manuscript. All authors edited the manuscript.

## Ethics declarations

Competing interests: The authors declare no competing interests.

