## Supplementary Information for "Key residue Unlocks Dual PIP_2_-Dependent and Independent Gating in G Protein-Gated Inwardly Rectifying Potassium Channels"

**Supplementary Figure 1. Molecular dynamics (MD) simulations of R92 and R92F GIRK2 channels.** (A) The root mean square deviation (RMSD) of the protein backbone (blue) and PIP_2_ (orange) over three independent MD simulations (MD1, MD2, and MD3) for R92 (i) and R92F (ii). (B) Root mean square fluctuation (RMSF) plots across the three simulations, comparing the flexibility of residues in R92 (i) and R92F (ii) channels. (C) Time-resolved analysis of the distance between the C-alpha carbon of R92 (i) or R92F (ii) and the 1’ PO_4_ of PIP₂ over three MD simulations.

**Supplementary Figure 2. Hydrogen bond analysis between PIP_2_ and R92 or R92F in MD simulations.** (A) The number of hydrogen bonds formed over time between the 1’-phosphate of PIP_2_ and R92 (pink) or R92F (blue) for 3 independent simulations (i-iii). (B) Probability of hydrogen bond formation between PIP_2_ and R92 (pink) or R92F (blue), mean and single points plotted.

**Supplementary Figure 3. Cryo-EM reconstruction of GIRK2 R92F.** Flowchart of image processing in cryoSPARC v3.3. Half-map of the final round of non-uniform refinement of State 1 and State 2 were sharpened by DeepEMhancer. Fourier Shell Correlation (FSC) of the local refinements for State 1 and State 2, with dotted lines indicating a threshold of 0.143, are shown. Local resolution plot of the full channels (State 1 and State 2) and angular distribution of final particles are shown.

**Supplementary Figure 4. Cryo-EM reconstruction of GIRK2 R92Q.** Flowchart of image processing in cryoSPARC v3.3. A half-map of the final round of non-uniform refinement of State 1 was sharpened by DeepEMhancer. Fourier Shell Correlation (FSC) of the local refinement for State 1 is shown, with dotted lines indicating a threshold of 0.143. Local resolution plot of the full channel (State 1) and angular distribution of final particles are shown.

**Supplementary Figure 5. Cryo-EM density.** Densities for protein segments of GIRK2 R92F in State 1.

**Supplementary Figure 6. Cryo-EM density.** Densities for protein segments of GIRK2 R92F in State 2.
