## Supplementary figures and images for "Key residue Unlocks Dual PIP_2_-Dependent and Independent Gating in G Protein-Gated Inwardly Rectifying Potassium Channels"

### Supplementary Figures

Figure S1

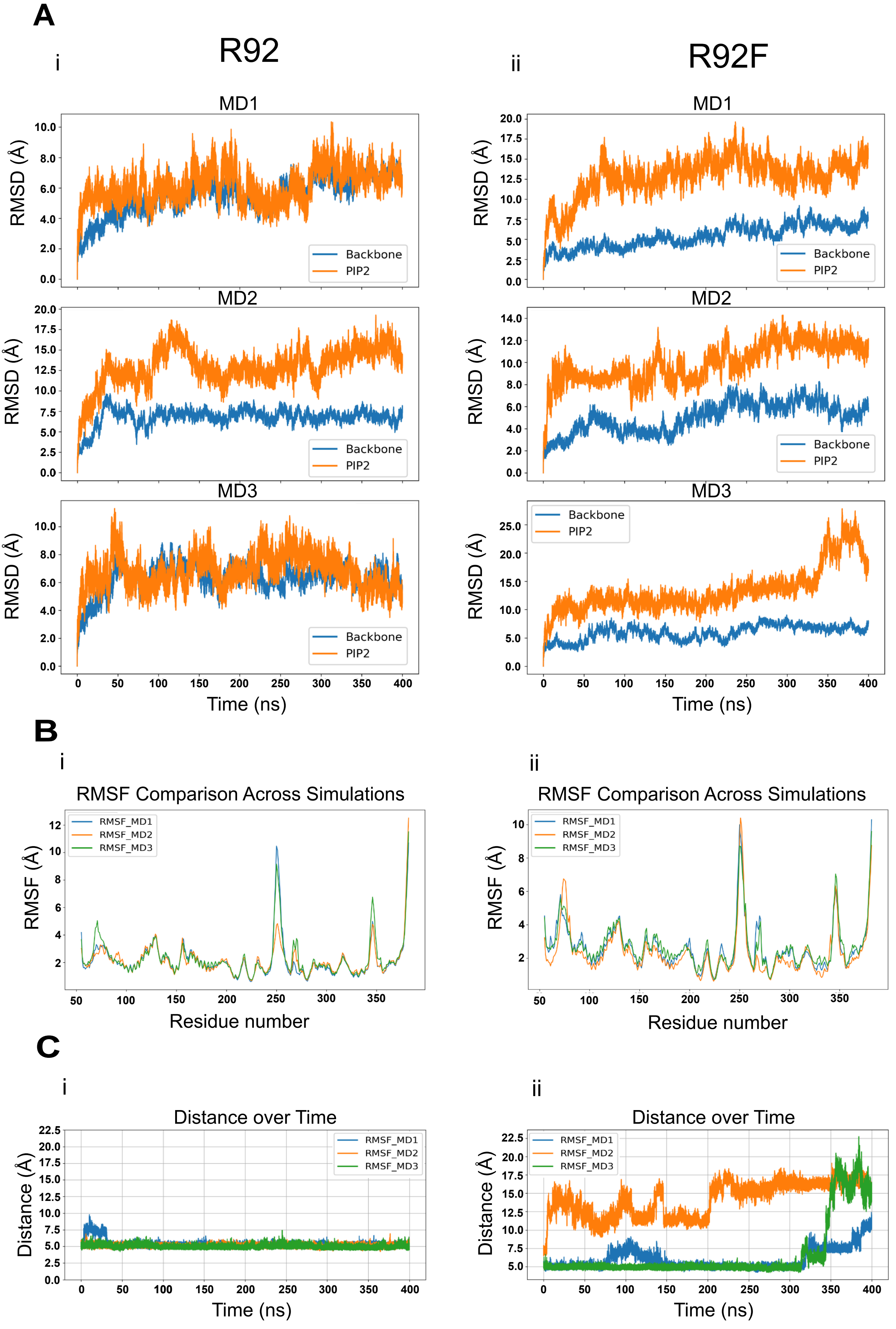


Figure S1

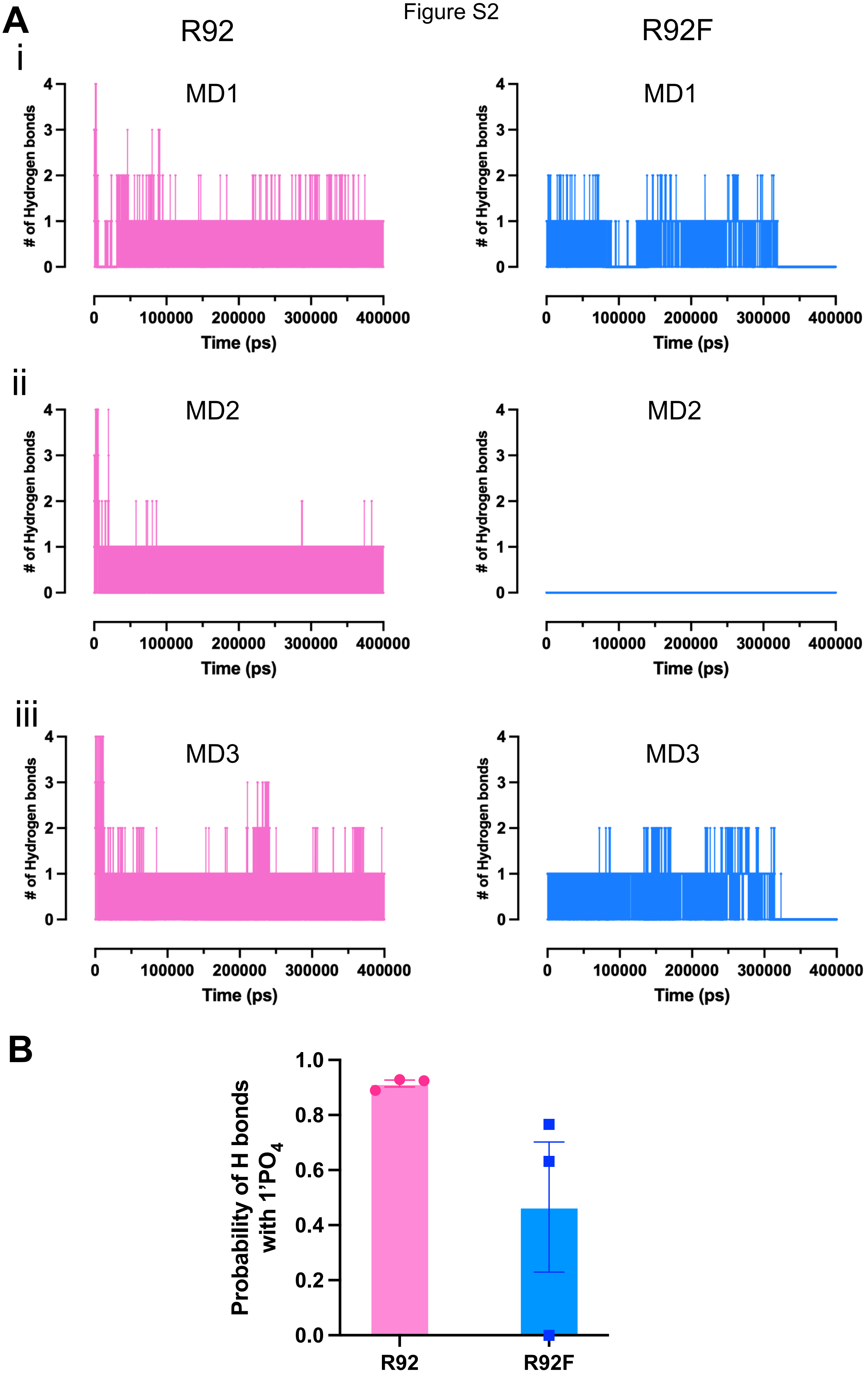

#### Figure S3

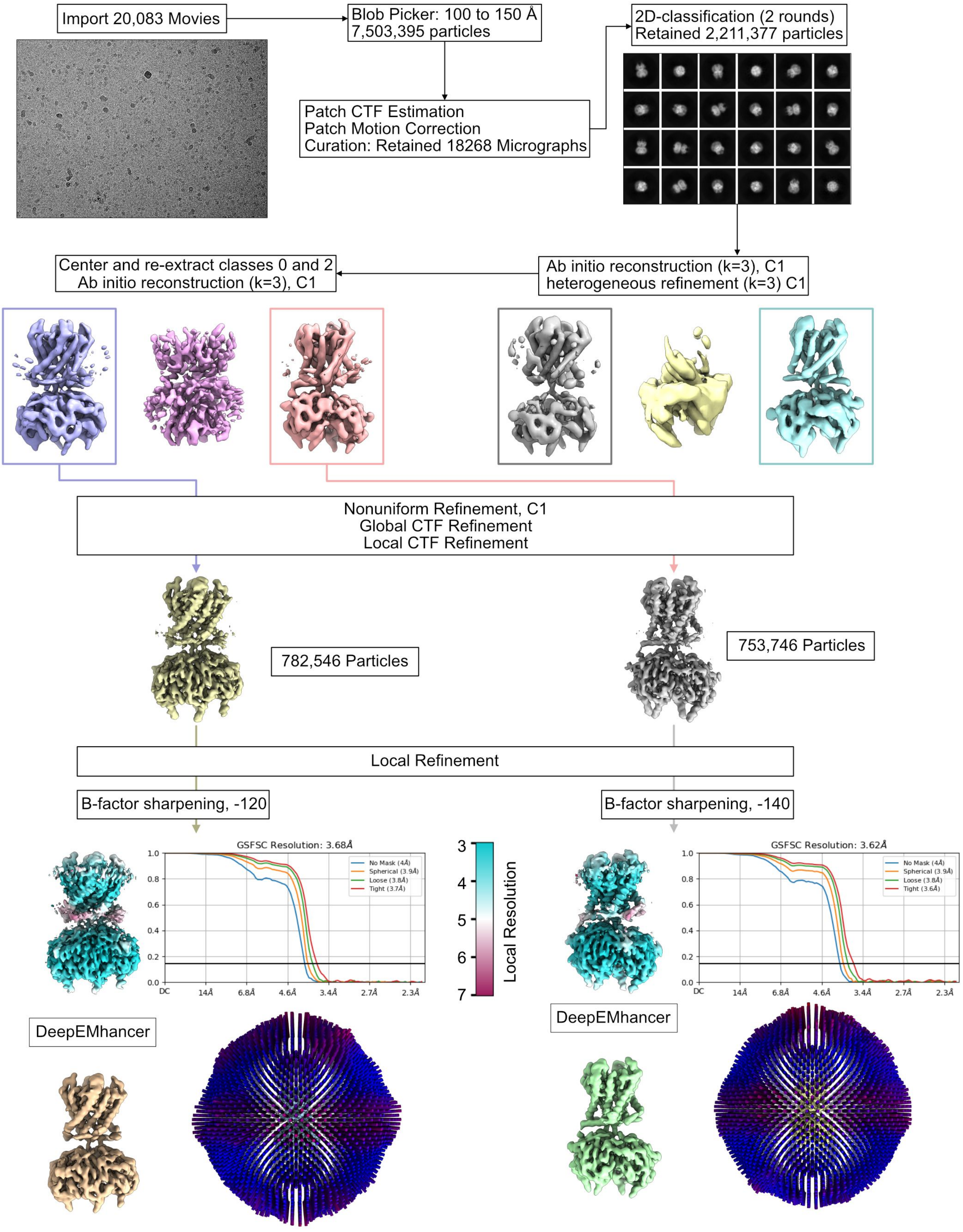


Figure S3

Figure S4

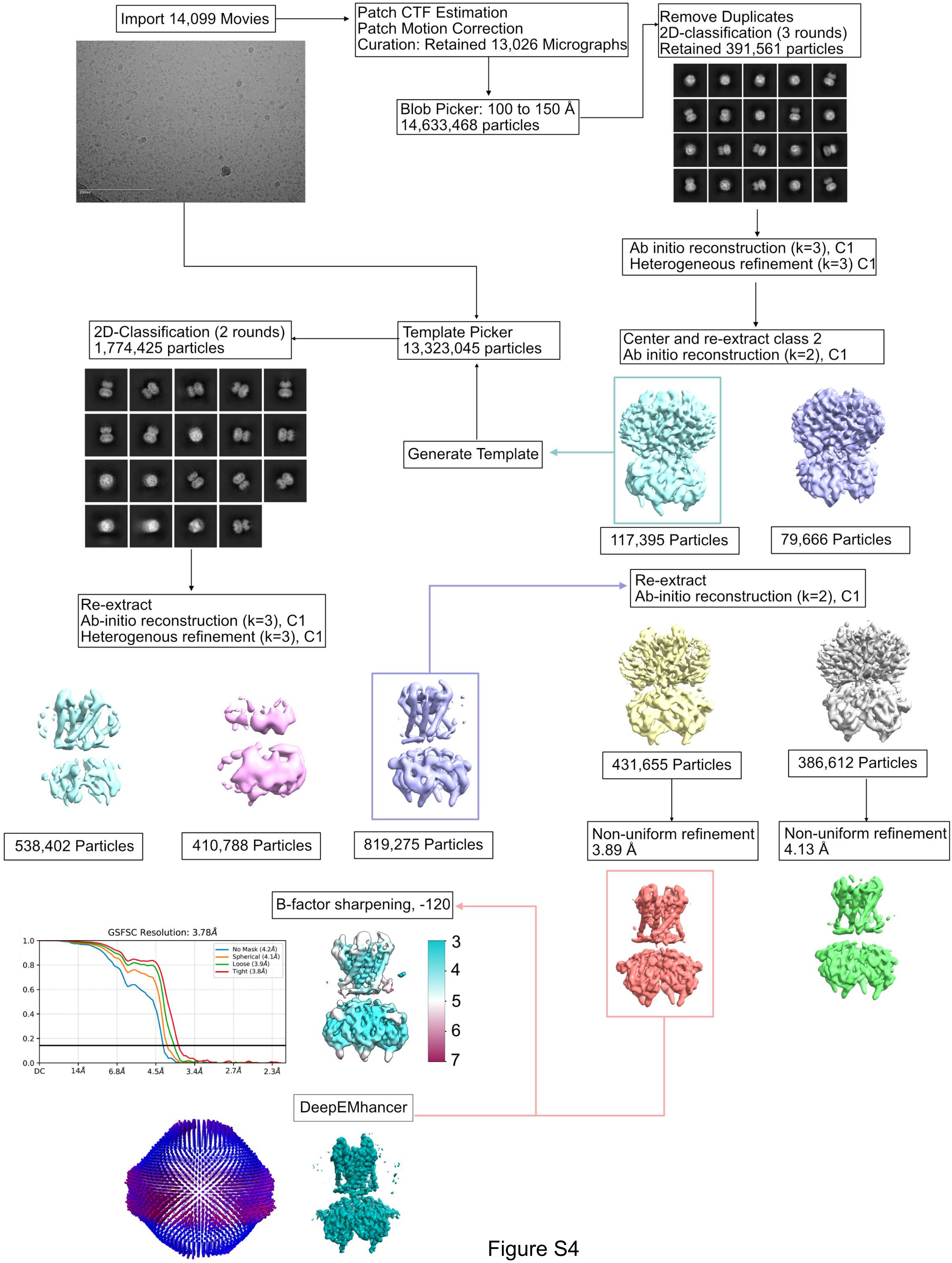


Figure S4

Figure S5

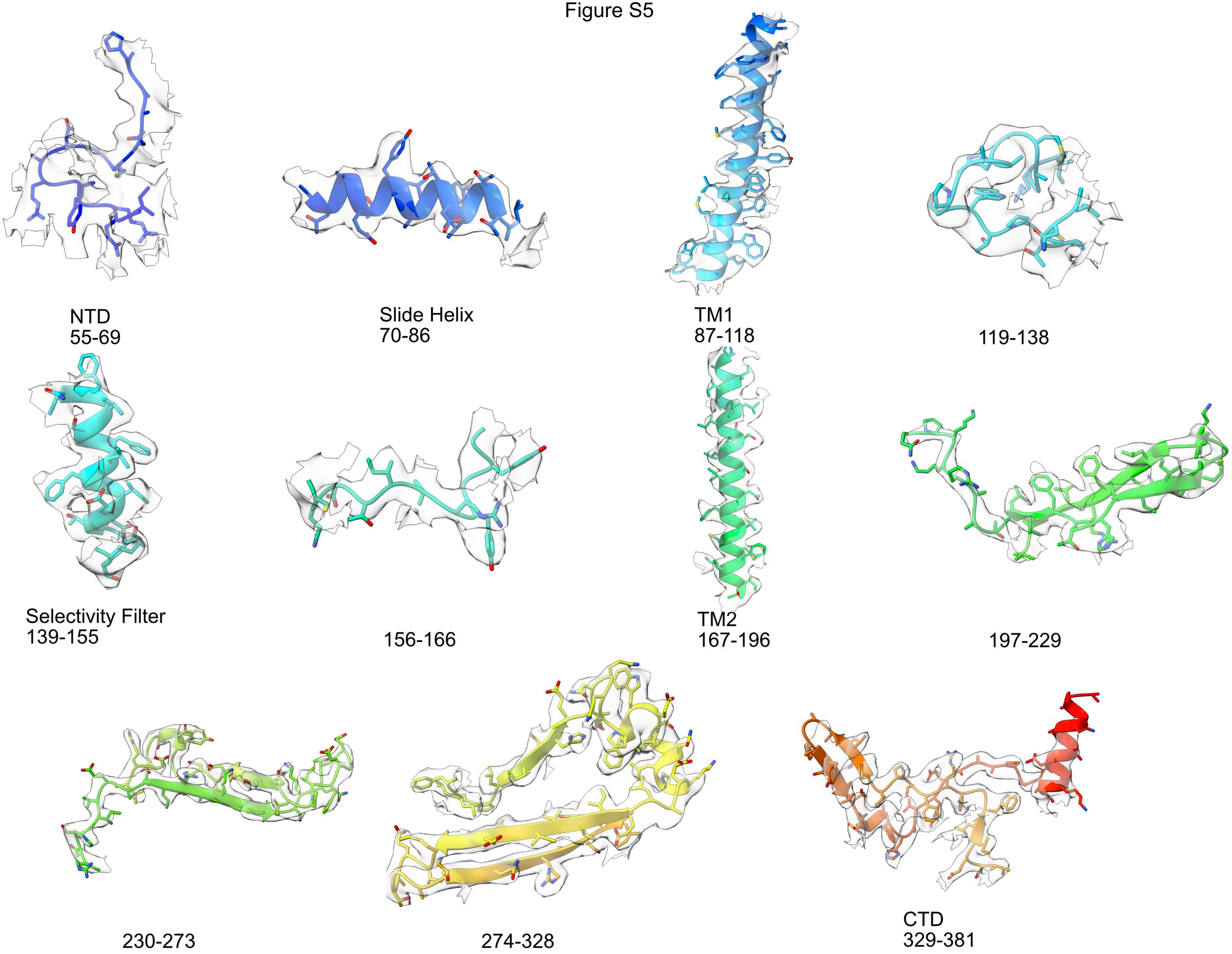


Figure S5

Figure S6

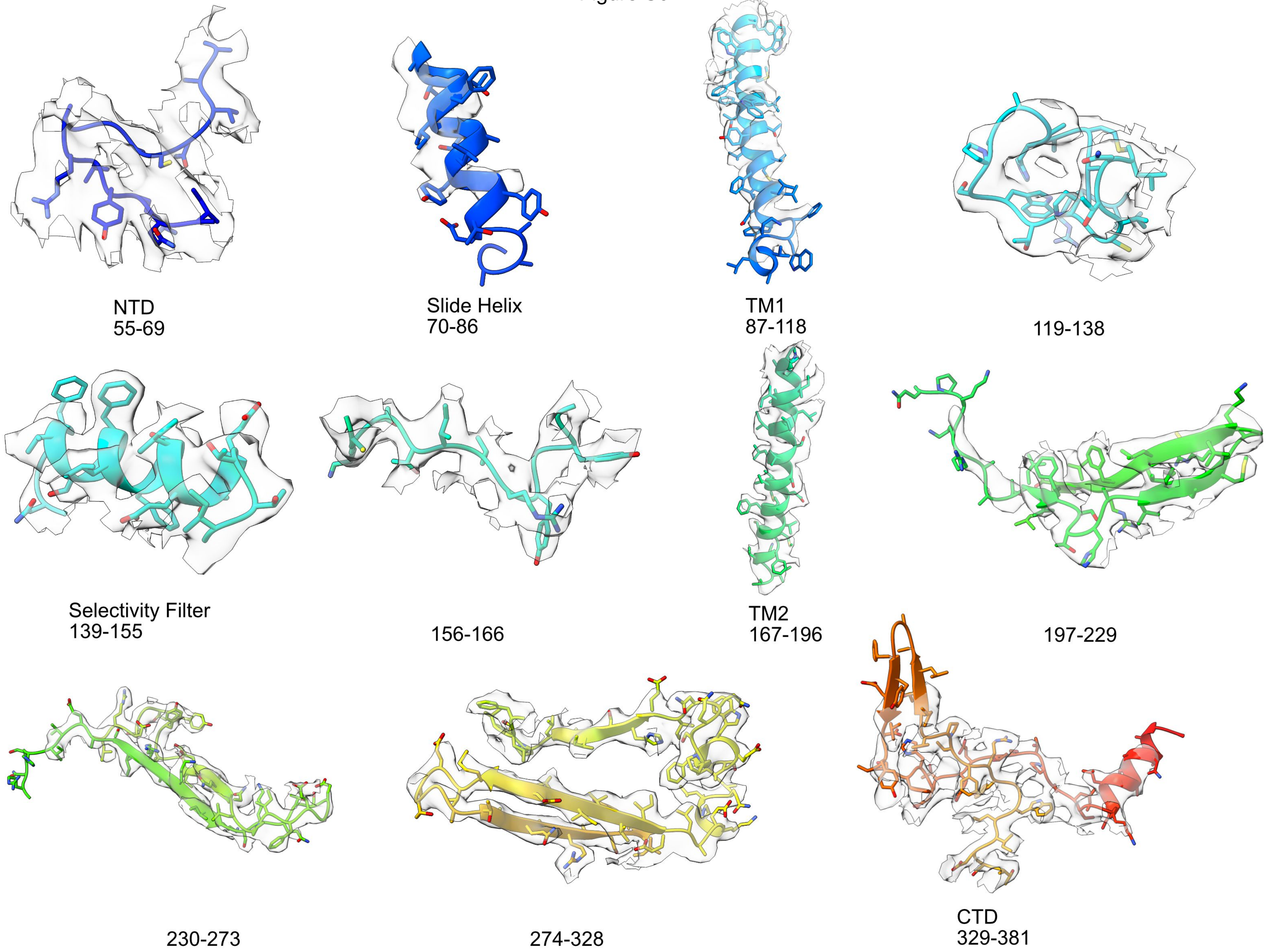


Figure S6
