## Supplementary Table 1 for "Key residue Unlocks Dual PIP_2_-Dependent and Independent Gating in G Protein-Gated Inwardly Rectifying Potassium Channels"

**Supplementary Table 1. Cryo EM data collection, refinement and validation statistics**

| | GIRK2 R92F State 1 (EMD-48268) (PDB 9MH9) | GIRK2 R92F State 2 (EMD-48267) (PDB 9MH8) | GIRK2 R92Q State 1 (EMD-48270) | GIRK2 R92Q State 2 (EMD-48269) |
|---|---|---|---|---|
| **Data collection and processing** | | | | |
| Magnification | 81,000x | 81,000x | 81,000x | 81,000x |
| Voltage (kV) | 300 | 300 | 300 | 300 |
| Electron exposure (e–/Å$^2$) | 53.92 | 53.92 | 51.73 | 51.73 |
| Defocus range (μm) | -2.7 to -0.6 | -2.7 to -0.6 | -3.6 to -0.4 | -3.6 to -0.4 |
| Pixel size (Å) | 1.069 | 1.069 | 1.058 | 1.058 |
| Symmetry imposed | C1 | C1 | C1 | C1 |
| Initial particle images (no.) | 698,029 | 936,163 | 1,774,425 | 1,774,425 |
| Final particle images (no.) | 88,923 | 122,422 | 431,655 | 386,612 |
| Map resolution (Å) | 3.16 | 3.58 | 3.89 | 4.13 |
| FSC threshold | 0.143 | 0.143 | 0.143 | 0.143 |
| Map resolution range (Å) | 3-6 | 3-6 | 3.5-7 | 4-7 |
| **Refinement** | | | | |
| Initial model used (PDB code) | 6XEU | 9MH9 | | |
| Model resolution (Å) | 3.20 | 3.20 | | |
| FSC threshold | 0.143 | 0.143 | | |
| Model resolution range (Å) | 2.9-3.4 | 3-6 | | |
| Map sharpening *B* factor (Å$^2$) | -120 | -120 | | |
| Model composition | | | | |
| Non-hydrogen atoms | 9764 | 9783 | | |
| Protein residues | 1277 | 1277 | | |
| Ligands | 0 | 0 | | |
| *B* factors (Å$^2$) | | | | |
| Protein | 88.95 | 162.54 | | |
| Ligand | N/A | N/A | | |
| R.m.s. deviations | | | | |
| Bond lengths (Å) | 0.004 (0) | 0.004 (0) | | |
| Bond angles (°) | 0.953 (0) | 0.984 (2) | | |
| Validation | | | | |
| MolProbity score | 1.87 | 1.83 | | |
| Clashscore | 8.64 | 9.80 | | |
| Poor rotamers (%) | 0.30 | 0.40 | | |
| Ramachandran plot | | | | |
| Favored (%) | 93.98 | 95.49 | | |
| Allowed (%) | 5.38 | 4.35 | | |
| Disallowed (%) | 0.63 | 0.16 | | |
